# ForSys: non-invasive stress inference from time-lapse microscopy

**DOI:** 10.1101/2024.05.28.595800

**Authors:** Augusto Borges, Jerónimo R. Miranda-Rodríguez, Alberto Sebastián Ceccarelli, Guilherme Ventura, Jakub Sedzinski, Hernán López-Schier, Osvaldo Chara

**Affiliations:** Unit Sensory Biology and Organogenesis, Helmholtz Zentrum München, Munich, Germany; Graduate School of Quantitative Biosciences, Ludwig Maximilian University, Munich, Germany; Instituto de Neurobiología, Universidad Nacional Autónoma de México (UNAM), Boulevard Juriquilla 3001, Juriquilla, México; Institute of Physics of Liquids and Biological Systems, University of La Plata, La Plata, Argentina; The Novo Nordisk Foundation Center for Stem Cell Medicine (reNEW), University of Copenhagen, Blegdamsvej 3B, 2200, Copenhagen, Denmark; Division of Science, New York University Abu Dhabi, Saadiyat Island, United Arab Emirates; School of Biosciences, University of Nottingham, Sutton Bonington Campus, Nottingham, LE12, UK; Instituto de Tecnología, Universidad Argentina de la Empresa, Buenos Aires, Argentina

## Abstract

During tissue development and regeneration, cells interpret and exert mechanical forces that are challenging to measure in vivo. Therefore, stress inference algorithms have emerged as powerful tools to estimate tissue stresses. However, how to incorporate tissue dynamics effectively into the inference remains elusive. Here, we present ForSys, a Python-based software that estimates intercellular stresses and intracellular pressures using time-lapse microscopy. We validated ForSys in silico and in vivo using the well-characterized mucociliary epithelium of the Xenopus embryo. We applied ForSys to study the migrating zebrafish lateral line primordium. We found that stress increases during cell rounding just before cell division and predicted the onset of epithelial rosettogenesis with high accuracy. Finally, we analyzed the development of the zebrafish neuromast and inferred mechanical asymmetries in a cell type-specific adhesion pattern. The versatility and simplicity of ForSys enhance the toolkit for studying spatiotemporal patterns of mechanical forces during tissue morphogenesis in vivo.

## Introduction

Recent advancements in experimental techniques have reignited interest in exploring the mechanical properties of biological tissues, commonly referred to as tissue rheology. These methods have facilitated precise and quantitative measurements of tissue mechanical parameters. For example, implanted deformable magnetic droplets have been used to determine the elastic properties along the zebrafish anteroposterior axis during body elongation ^1,2^ and presomitic mesoderm differentiation ^3^. Similarly, the application of optical traps has enabled controlled deformation of cell membranes, thereby facilitating the study of viscoelastic properties during *Drosophila* development ^4^. Laser ablation experiments have also been employed in various systems to probe cortical tension by measuring the recoil of cell junctions upon laser cutting ^5–7^. Despite their importance, these experimental methods often come with significant drawbacks. They can be costly and necessitate specialized equipment, posing implementation challenges for many researchers. Moreover, these techniques might not be conducive to long-term imaging, potentially disrupting the normal development and, in some cases, leading to the destruction of the sample. Hence, there is a pressing need for alternative approaches to overcome these limitations while still delivering accurate and non-invasive measurements of tissue rheology.

Computational methods offer a promising solution, enabling the cost-effective and straightforward implementation of tissue mechanical characterization *in vivo*^8^. Inference techniques have emerged as powerful tools in tissue rheology, utilizing readily available microscopy images to infer the effective stress of a system based on the geometry of the cellular junctions. A key aspect of this approach centers on tricellular junctions (TJ), where three cells converge ^9^. The underlying framework relies on one major assumption: mechanical equilibrium is maintained at each TJ. The strength of these models lies in their simplicity, reducing the estimation of intercellular stresses to the solution of an overdetermined system of linear equations ^10,11^. One of the first implementations of the force-inference approach is CellFIT ^10^, which enables the estimation of stresses from microscopy images. While CellFIT provides accurate estimates in static tissues, its applicability to dynamic tissues is limited. Although recent techniques using time series data offer improvements ^12^, a computational tool capable of dynamic stress inference has been lacking.

Here, we introduce ForSys, an open-source Python-based inference algorithm specifically developed to tackle the complexities of dynamical stress inference from time series experiments. ForSys utilizes the local velocity of cell junctions to extract the spatiotemporal stress distribution *in vivo*, providing accurate estimations of a tissue’s mechanical state.

## Results

### ForSys: a Python-based open-source software to infer mechanical stress in tissues

ForSys enables the inference of intercellular mechanical stress and intracellular pressure of tissues. It takes the two-dimensional (2D) segmentation of an image, which delineates cell outlines, as its input. It then conceptualizes the entire tissue as a polygonal structure. In this structure, each polygon represents a cell, with edges connecting vertices.

ForSys operates in two distinct modes contingent upon the input (Fig. 1). When supplied with a singular segmentation of a static image, the software engages its Static mode (Fig. 1A) (see “Statical stress inference” in the Materials and Methods section). In this mode, a stress inference is applied to a single image. Conversely, if the input comprises the segmentation of a time series dataset, ForSys presents the option to function in its Dynamic mode (see “The dynamic inference case” in the Materials and Methods section) (Fig. 1B). This mode involves the extraction of temporal trajectories for vertices from the microscopy time series, thereby incorporating corresponding vertex velocities to refine stress inference.

**Figure 1.**
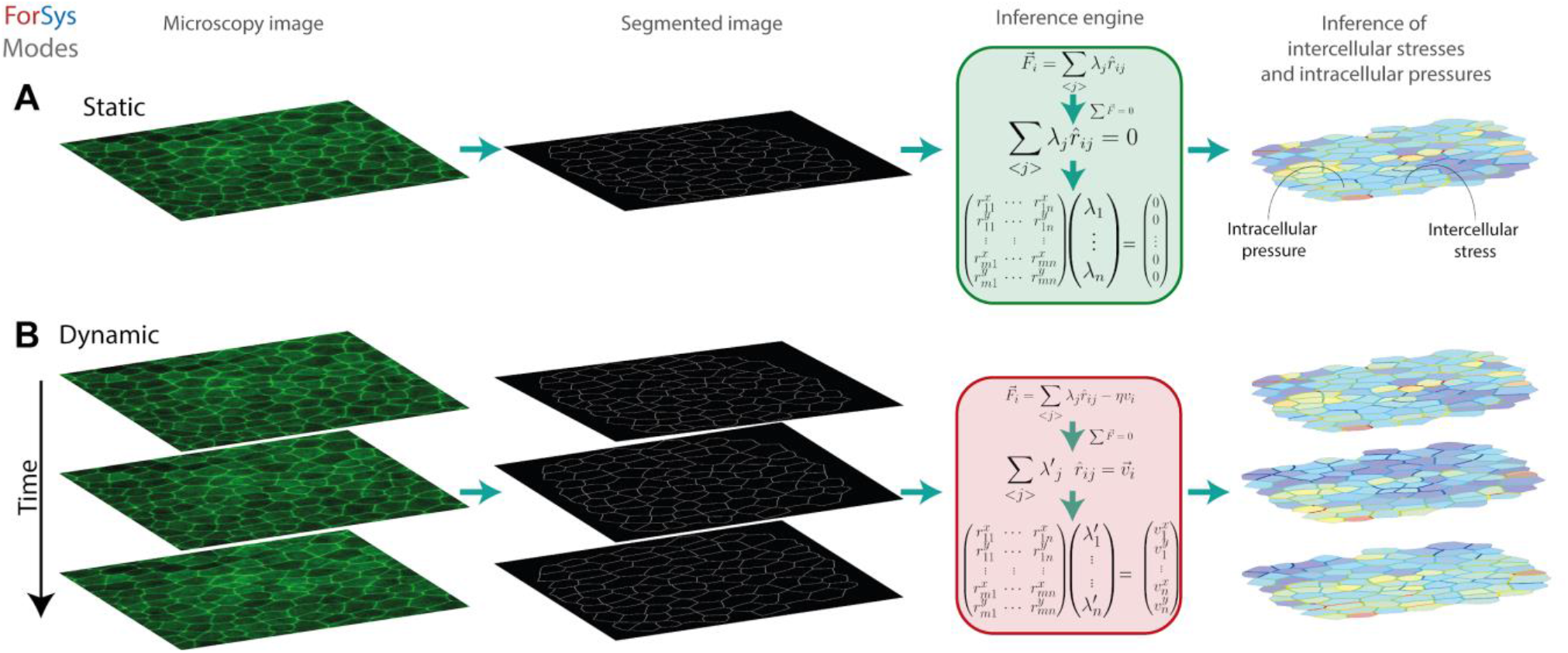
Force inference modalities of ForSys. **(A)** The static inference is performed on a microscopy image by creating a skeletonized tissue representation. Then, ForSys reads it and builds the system of equations according to the geometrical properties of the tissue, assuming that each vertex in the tissue is in mechanical equilibrium. Lastly, the system is solved, and the intracellular pressures and intercellular stresses are inferred. **(B)** Similarly, the dynamical inference uses a time series of images to add dynamical information to the system of equations used in the static case, by assuming an overdamped regime. A time mesh is generated from the succession of microscopy images, and pivot vertices are tracked through time. These are vertices at which three or more edges meet. Then, the velocity of these vertices from frame to frame is used to modify the system of equations, allowing non-static tissues to be analyzed by stress inference.

### ForSys infers *in silico* stresses accurately in static equilibrium

To assess ForSys’s performance against existing tools, we utilized as a ground truth simulations generated by a vertex model implemented in Surface Evolver via seapipy, analyzed it using our software, and compared the results with outputs from previously published methods, focusing specifically on CellFIT ^10^ and DLITE ^12^. Given that both tools yield similar results (Extended data figure 3A and 3B), we opted for DLITE implementation due to its open-source nature, enabling a direct comparison with tissue stresses extracted from Surface Evolver outputs.

We selected the final time-point (*t* = 24) of simulations generated from four different conditions to compare the ground truth from the Surface Evolver output (Fig. 2A), DLITE’s estimation (Fig. 2B), and ForSys in its Static modality (Fig. 2C). In all cases, the predicted intercellular stresses and intracellular pressures closely matched the ground truth. Moreover, both stress inference methods exhibited a high degree of accuracy and precision, as reflected by a low Mean Absolute Percentage Error (MAPE) (<10%) (Fig 2D) and a high saturated score function (∼30) (Figure 2E and Extended Data Figure 3C). Importantly, ForSys showed a significantly lower MAPE (*p* = 1e-05; *p* = 1e-8; *p* = 0.01, for the x-furrow, y-furrow, and circular furrow, respectively), higher saturated score (*p* < 0.001; *p* < 1e-4; *p* < 0.01 for the x-furrow, y-furrow, and circular furrow, respectively) and smaller interquartile range than DLITE, for all cases except the random tensions (See the “Statistical estimator” section in Materials and Methods for details).

**Figure 2.**
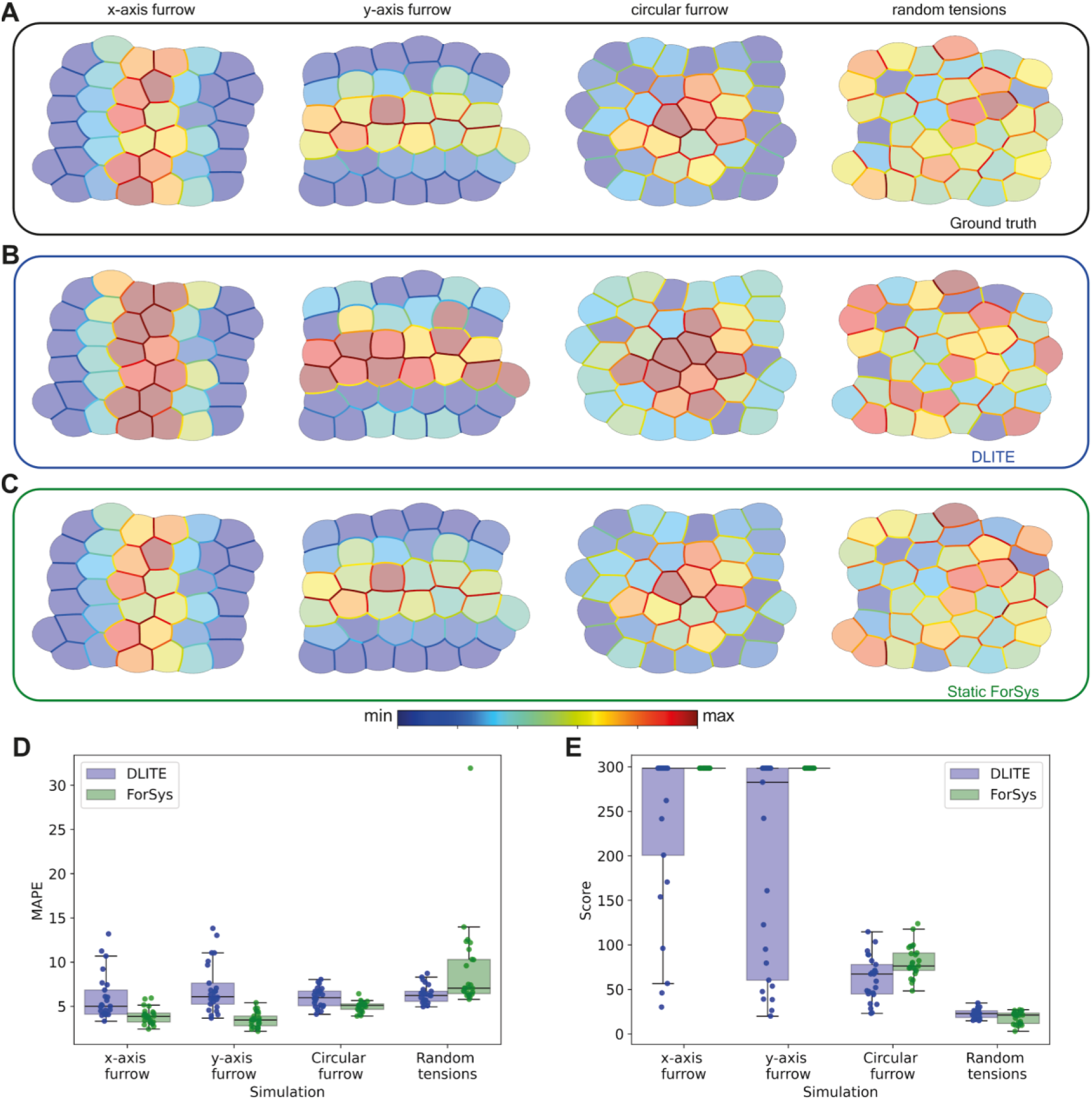
*In silico* validation of ForSys for tissues in static equilibrium. Four different conditions were generated with seapipy to benchmark ForSys under the static equilibrium condition. Each column shows a representative replicate per condition at the final frame (t=24). The ground truth **(A)** can be compared to the values for the DLITE predictions **(B)** and the Static ForSys **(C)**. The three rows shown correspond to the final frame of the simulation. The color bar above the last two panels shows the order of the colormap for both the stresses and the pressures. Pressures in the cells are represented with transparency for improved visualization. The mean absolute percentage error (MAPE) **(D)** and the saturated score function **(E)** for all simulations (N = 25) are represented in two boxplots, DLITE and Static inference with ForSys, paired by condition. (see Materials and Methods “Evaluating goodness of fit) Dots show the result for individual repetitions.

These results indicate that ForSys’s static modality yields higher accuracy and precision estimations than DLITE while effectively capturing the *in silico*-generated ground truth spatial distributions in static equilibrium. Consequently, only ForSys in its Static modality will be used hereafter for comparison with a static solution.

### ForSys stress inference in dynamical tissues outperforms static methods *in silico*

With the aim of inferring stress in dynamic tissue, we assumed that the tissue goes through a succession of quasistatic states in an overdamped regime^13^, consistent with a viscoelastic response of the cell junctions to the deformations created by the forces acting on them^14,15^. Consequently, we incorporated a viscous term proportional to the velocity of the corresponding vertex in each junction’s equation. Importantly, these velocities are not unknown: ForSys estimates them using the spatial coordinates of the vertices tracked over time. In ForSys, we call this modality of stress inference Dynamic.

Dynamic inference depends on a dimensionless parameter proportional to the reciprocal of the Weissenberg number^16–18^ (see a detailed description in the “The dynamic inference case” section of Materials and Methods). Thus, we fitted this parameter and found its optimal value for each of the examples. Our results indicate that the best dynamic results are obtained with a scale parameter of about 0.1 (see Extended Data Fig. 5 and the Materials and Methods section “Determination of the scale parameter”).

Under our prescribed conditions (Figure 3A), ForSys in its dynamical modality (Fig. 3C) outperforms static inference (Figure 3B), accurately reproducing stress and pressure distributions akin to the ground truth. Furthermore, our results indicate that dynamic modality improves static modality accuracy and precision, as indicated by MAPE (p < 1e-09; p < 1e-9; p < 1e-7, for the x-furrow, y-furrow, and circular furrow, respectively) (Figure 3D) and the saturated score function (p < 1e-08; p < 1e-9; p < 1e-9; p = 0.03, for the x-furrow, y-furrow, circular furrow, and random, respectively) (Figure 3E) (See the “Statistical estimator” section in Materials and Methods for details).

**Figure 3.**
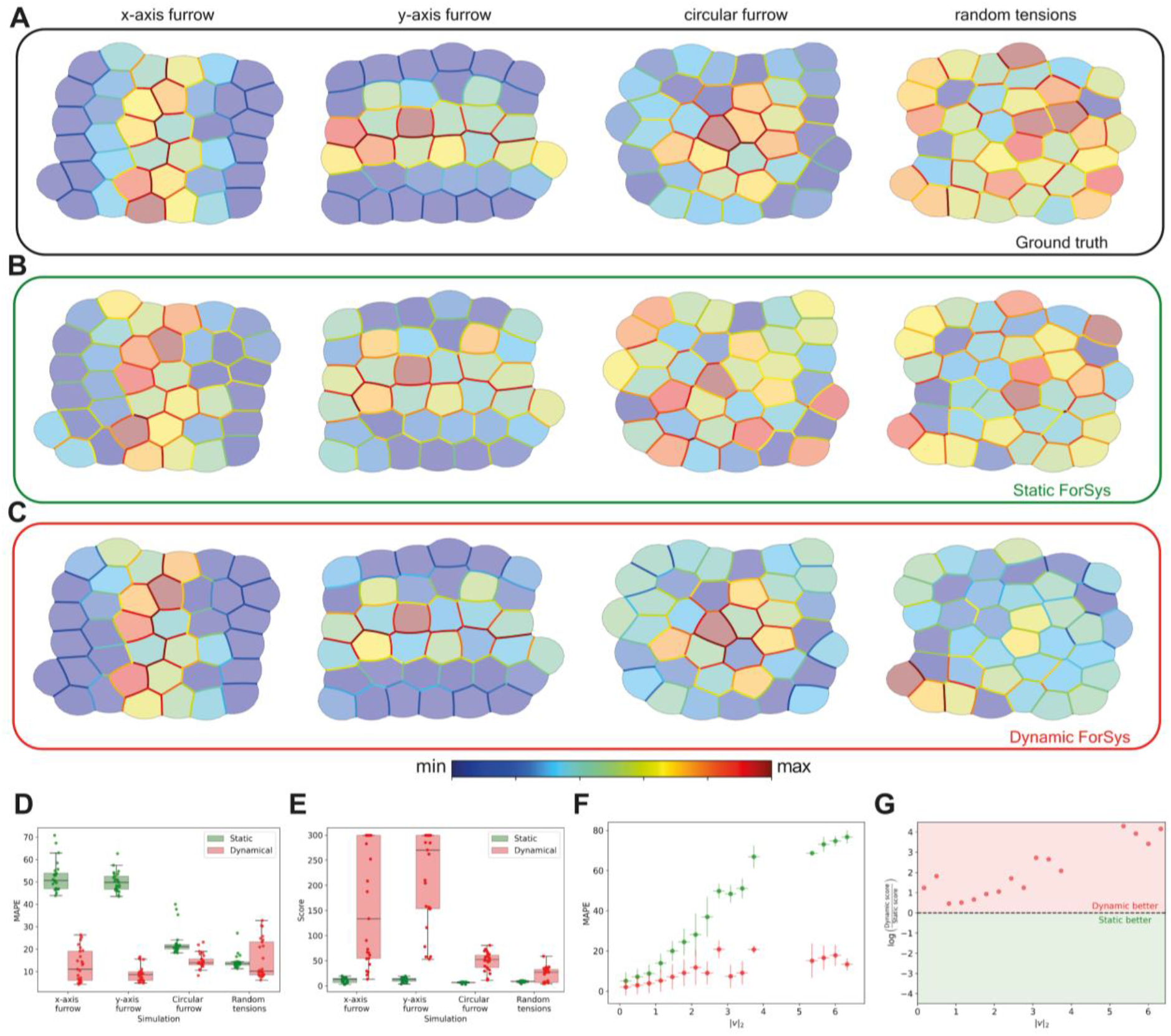
*In silico* validation of ForSys for tissues in dynamical equilibrium. We generated four examples with seapipy to test dynamical equilibrium conditions. Each column shows a representative repetition per example. The first row **(A)** shows the ground truth values for the stress and the pressures, the static inference made by ForSys is in the second row **(B)**, and the dynamical ForSys inference is in **(C)**. We show each example at one time point after the system’s tensions changed. The color bar above the last two panels shows the order of the colormap for both the stresses and the pressures. The mean absolute percentage error **(D)** and the saturated score function value **(E)** for all simulations are represented in two boxplots, Static and Dynamical inference, paired by condition. Dots show the result for individual repetitions. **(F)** Dependence of the MAPE with the velocity |*v*|_*2*_. The scattered dots are the median for all experiments with a velocity corresponding to the current |*v*|_*2*_ bin. Error bars in the y-axis are one standard deviation, and error bars in the x-axis represent the size of the velocity bin. **(G)** Dynamic to static score function ratio 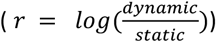 as a function of the |*v*|_*2*_ bin. A ratio bigger than zero shows that the dynamic solutions performed better (Red zone), and a negative value (Green Zone) favors the static solution. The black dashed line at y = 1 separates both zones. All velocity bins favor the dynamic solution.

Interestingly, accuracy and precision (estimated with the MAPE and the saturated score function) of stress inference in each ForSys modality are damped by the increases of TJ local movements, here reflected in the norm of the velocities vector (|*v*|_*2*_) (Figure 3F and G). Notably, the dynamic modality outperforms the static one for all TJ velocities, as observed by the time evolution of MAPE (Figure 3F). This can be evidenced through the ratio between dynamic and static scores (Figure 3G), where values greater than one mean that the dynamic modality outperforms the static one. The outperformance of the dynamical modality is clearer for higher TJ velocities (Figure 3F and Figure 3G and Extended Data Fig. 4). Thus, ForSys, in its dynamic modality, can retain a better approximation due to its use of the vertices’ velocity, *i*.*e*., future positions, to estimate the stress.

In this section, we have demonstrated through *in silico* validation that the dynamic modality of ForSys outperforms other methods in accurately inferring stresses in remodeling tissues.

### ForSys validation *in vivo* using the mucociliary epithelium of *Xenopus* embryos

To validate ForSys in a biological setting, we used published data from the mucociliary epithelium in *Xenopus* embryos (Fig. 4A) ^19^. We quantified myosin II intensity using a non-muscle myosin II A-specific intrabody (SF9-3xGFP, for simplicity referred to as myosin II), which has been previously used as a proxy for active myosin II ^20,21^. We segmented the microscopy images using Epyseg ^22^ (see Materials and Methods “SF9 myosin II sensor intensity measurements” for details) and compared myosin II measurements with the stress values inferred by ForSys.

**Figure 4.**
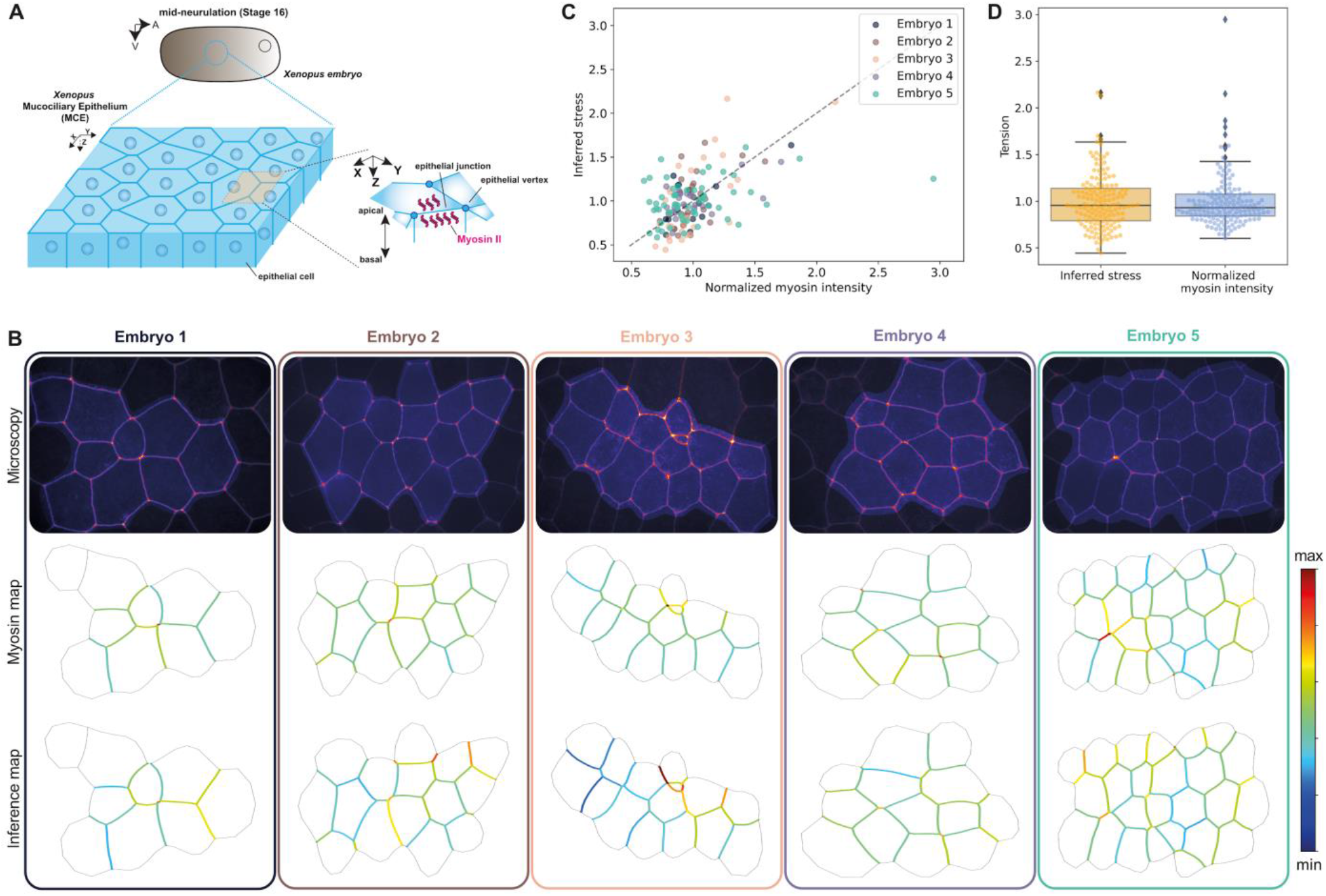
Comparison of ForSys-derived stress with myosin II measurements in the Xenopus embryo mucociliary epithelium. **(A)** Scheme of the Xenopus embryo and position of the mucociliary epithelium. **(B)** Five examples of inference in Xenopus embryos. The microscopy image is shown alongside the myosin intensity map and the ForSys inference result. The color code in the maps represents the myosin sensor intensity and the stress prediction. The scale was saturated at tension values of two. The highlighted region in the microscopies shows the area that was analyzed. **(C)** Relationship between myosin sensor intensity and stress inferred for the five examples. Each scatter point shows the value for a particular membrane in that example. The dashed black line represents the y=x line. Each color coincides with the rounded rectangle around the embryo and its font color in panel (B). The average Pearson correlation coefficient is R=0.56 ± *0*.*11;* (*m*ean±std) **(D)** Quantification of stresses and myosin sensor intensity for the five examples. Inferred stresses and myosin intensities are not significantly different from each other (p=0.76; Mann-Whitney U test; N=154).

As in previous sections we qualitatively compared the derived stress distribution maps with the ground truths, here given by the normalized myosin II sensor intensity (Figure 4B). We observed good qualitative agreement between inferred stress and myosin intensity, with regions of higher myosin fluorescence corresponding to higher inferred stress, most noticeable in Embryo 3 and Embryo 5 of Figure 4B. In contrast, in Embryo 4 of the same panel, ForSys can reproduce a more homogenous distribution along the tissue. On a quantitative level we found that ForSys predictions are moderately correlated with the myosin measurements for each embryo (R=0.56 ± 0.11; mean ± *std*) (*Fi*gure 4C). In addition, ForSys stresses predictions have a MAPE value of (21 ± 5)% (mean ± std). *Ov*erall, the distributions of myosin intensity and inferred stress are qualitatively similar and not significantly different (Figure 4D, p=0.76; Mann-Whitney U test; N=154).

Consequently, ForSys accurately and precisely infers the stresses present in the mucociliary epithelium of the Xenopus embryo, as measured by the fluorescence of the myosin II sensor.

### Dynamic stress inference of collective cell behavior in zebrafish

We sought to explore ForSys inferences in an *in vivo* model that mixes TJs with low and high motility. To this end, we turned to two morphogenetic processes that occur during the development and homeostasis of the zebrafish lateral line, a mechanosensory organ formed by a collection of discrete organs called neuromasts.

We first applied ForSys to an *in vivo* model of collective cell morphogenetic behavior leading to the formation of epithelial rosettes in the lateral line primordium of developing zebrafish. The primordium is a collection of just over 100 cells that move collectively from head-to-tail of the fish embryo (Fig. 5A). During migration, groups of approximately 25 trailing cells periodically detach from the primordium, sequentially giving rise to individual neuromasts that are then deposited at semi-regular pace ^23^. Although the lateral line primordium has been extensively characterized genetically ^24,25^, the mechanical forces present during migration and rosettogenesis remain unknown.

**Figure 5.**
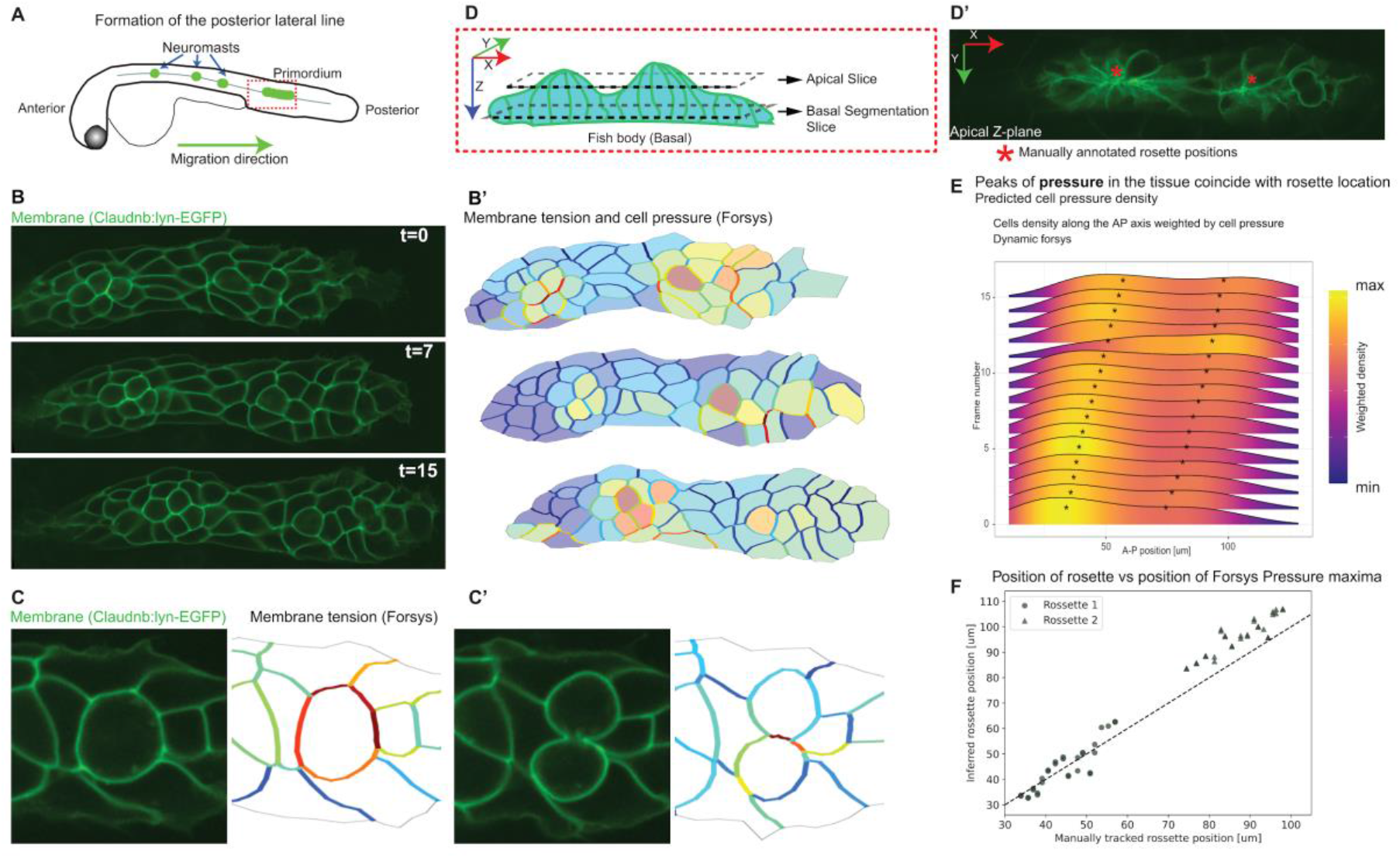
ForSys inference of a moving epithelium in the zebrafish lateral line at 2 dpf. **(A)** Schematic of the biological model. The neuromasts of the posterior lateral line are formed by detaching from a primordium that migrates from the anterior to the posterior of the fish. **(B)** Frames 0, 7, and 15 of the primordium migration in which cell membranes are fluorescently marked with Claudnb:lyn-EGFP. **(B’)** The membrane signal is used for segmentation, which ForSys can use to predict cell membrane tension and intracellular pressure. **(C, C’)** Consecutive planes show cell division. The membrane tension in the cell just about to divide is considerably higher than the surrounding membranes. After division, the dividing membrane retains a high tension. **(D, D’)** Schematic of the primordium orientation and the position of the optical planes. Constriction of the cell membranes in rosettes is evident in the apical plane. The asterisks show the anteroposterior location of rosettes. The cell segmentation was done on a Z-plane at a more basal plane **(E)** Ridgeline plots of Cell densities along the anteroposterior axis throughout 16 frames for a representative primordium. Time goes from bottom to top. The direction of primordium migration is to the right. The asterisks show the positions of the manually annotated rosettes. **(F)** Anteroposterior position of the manually tracked rosette against the inferred position by taking the local maxima of the density of pressure values from (E). The diagonal line marks y=x as a reference for comparing predicted and manually annotated values.

Therefore, we decided to use ForSys in its Dynamic modality to analyze time lapse data of migrating primordia, whose cells’ plasma membranes were fluorescently labeled with EGFP. Migrating primordia were followed for 30 minutes with a temporal resolution of two minutes (Supplementary video 1)(Fig. 5B and 5B’).

ForSys predicted the mitotic division of primordial cells by revealing high stress in the pre-dividing cellular membrane relative to the membrane of the non-dividing surrounding cells (Fig. 5C). The stresses remain partially conserved after division, mainly in the cell membrane separating the resulting cell siblings (Fig. 5C’).

We then applied ForSys to predict stress tissue-wide. Apical constrictions of epithelializing cells are mechanistically associated with the formation of the rosettes that preempt neuromast morphogenesis ^26^. The apical constriction is readily detectable by morphology when looking at the apical plane of the primordium (Fig. 5D and 5D’)^27^. The relationship between apical constrictions and forces in more basal planes of the cells and how they relate to rosettogenesis remain undefined.

To begin to address this possible relationship, we used time series data and aggregated the position of the cells along the anteroposterior axis of the primordium by kernel density estimation. We weighed each cell using the intracellular pressure inferred from ForSys, which results in a smoothed curve estimating intracellular pressure along the migration axis (Fig 5E). This analysis showed that the anteroposterior positions of the rosettes, manually annotated by looking at apical constriction (Asterisks in Fig. 5D’ and 5E), correlate with the predicted zones of high intracellular pressure inferred by ForSys. The closeness between the predicted pressure maxima and the manually annotated rosette formation indicates a high correlation between these two quantities during primordium migration (R=0.99, p<1e-51, N=61; for rosettes 1 and 2 combined) (Figure 5F). Encouraged by our previous results, we next analyzed mature neuromats. The center of this organ is occupied by mechanosensory hair cells, which are surrounded by non-sensory supporting cells (Figure 6A) ^28^. We used a plasma membrane marker to define cells, which were segmented using ilastik ^29^ and epyseg ^22^ (Figure 6B). Then, we used the Dynamical modality of ForSys to estimate the stress at each membrane (Figure 6B’) and found that membranes belonging to hair cells have higher stress on average. Homotypic interfaces between hair cells have the highest stress (p < 1e-7 vs. hair cell-supporting cell interfaces). On the other hand, homotypic contacts between supporting cells have the lowest stress (p<0.006 vs. hair cell-supporting cell interfaces) (Figure 6C).

**Figure 6.**
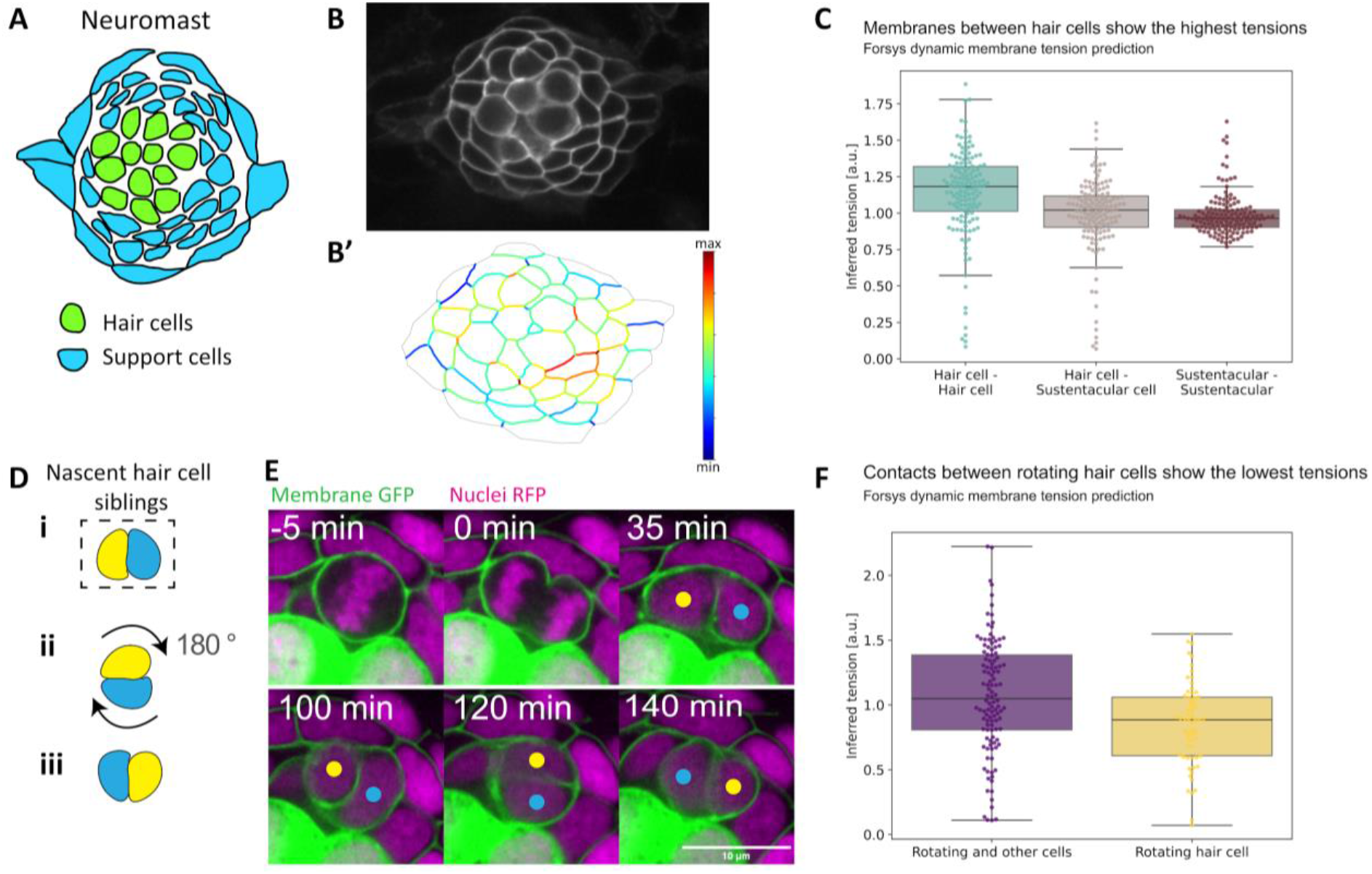
*in vivo* ForSys inference in an epithelium with rotating cells. **(A)** Schematic of cell composition in a zebrafish lateral line neuromast. Sensory hair cells are located in the center and are surrounded by support cells. **(B)** Image of a neuromast whose cells can be tracked by membrane-tethered EGFP. **(B’)** ForSys tension inference after membrane segmentation. **(C)** The tension inferred for membranes is classified by the type of cell-cell contact. The homotypic contacts between hair cells show the highest predicted tension, while the homotypic contacts between support cells show the lowest on average. Each data point is the mean of the predicted tension values for each membrane type in one frame. The frames come from N=7 time-lapse experiments. **(D)** Schematic of the planar cell inversions occurring in 50% of the nascent hair cell pairs: sibling hair cells perform a 180° rotation to exchange positions along the anterior-posterior axis. **(E)** Time-lapse frames showing the in vivo rotation process: around 100 minutes after mitosis, the nascent hair cells exchange anteroposterior positions by rotating in the epithelial plane. The sibling cells remain attached to each other during the rotation, while the surrounding cells do not actively participate in the movement. **(F)** Homotypic tensions between the young rotating hair cells are significantly lower than their contacts with the surrounding cells.

We then focused on a still-puzzling process called planar cell inversion (PCI) ^30,31^. PCI occurs when supporting cells give rise to hair-cell progenitors, which divide once to generate a pair of hair cells. Approximately half of the resulting nascent hair-cell pairs undergo a 180º rotation around their geometric center ^30,31^ (Figure 6D and 6E). The mechanical forces occurring during cell-pair inversions are not known. Therefore, we focused our analysis on the homotypic junctions between the sibling hair cells and compared them to those with the surrounding supporting cells. This allowed us to test how the stress differs between the different cell types. We found that the stress in the membranes juxtaposing the rotating hair-cell pair is significantly smaller than that between hair cells and the adjacent supporting cells (p<0.0005) (Figure 6F). Because tension and adhesion are generally inversely related, PCI could be characterized by a strong adhesion within the rotating cell pair and weaker adhesion with the surrounding cells. This result suggests a cell-type and cell-state-specific adhesion pattern that underlies contact remodeling necessary for coordinated cell-pair rotations. Taken together, these results show that ForSys’s dynamical implementation predicts high stresses before cell division in a migrating tissue. They also revealed that rosette formation could be prefigured by mechanical rosettogenesis changes in the cells, which allows the inference of apical constrictions during rosettogenesis using information from basal planes.

## Discussion

Here, we introduce ForSys, a new software that statically and dynamically infers stresses without disrupting biological tissues. Traditional inference methods rely on geometrical information to calculate the relationship among the stresses acting on cell membranes in a static image. However, these methods generally lack the dynamical component present in a time-series microscopy. ForSys extends the applicability of inference techniques by enabling dynamic stress inference in cell membranes when tissues are in motion.

We validated our software in its static and dynamic modalities with different *in silico* spatial patterns of tissue stresses using a cell-based computational model implemented in Surface Evolver ^32^, which we integrated into a Python package called seapipy ^33^. Our results show that ForSys can recover the ground truth in its static and dynamic modalities. Significantly, the dynamic modality improves the accuracy of the static modality. Unlike static inference, characterized by a unique scale contained in the stresses to be inferred, dynamic inference adds a viscous term proportional to the nodes’ velocities, which introduces an additional scale to the problem. The dynamic inference can thus be reformulated in terms of the Weissenberg number ^16–18^. The optimal value found for this number *in silico* indicates that elastic forces are an order of magnitude larger than viscous forces. Strikingly, dynamic inference outperforms static inference even when elastic forces dominate over the viscous forces, pointing to a wide applicability of the dynamic modality.

We then validated ForSys in the Xenopus embryonic mucociliary epithelium. We found a positive correlation between the inferred stress and cortical stress that was indirectly measured using variations in the intensity of myosin II. As the embryonic mucociliary epithelium progresses over several hours, continuous, direct probing of the mechanical forces is extremely laborious, likely to interfere with tissue development, and hardly compatible with single-cell resolution measurements. Therefore, using ForSys for non-invasive mapping of mechanical forces at the scale of an entire tissue across time could pave the way for a more comprehensive understanding of the mechanical forces that drive tissue development.

We further demonstrated the power of ForSys by studying two aspects of organ development and homeostasis using the neuromasts of the lateral line in zebrafish embryos. Specifically, we addressed two processes that involve a complex collective cell behavior. First, we applied Dynamical ForSys to the migrating lateral-line primordium. Although this process has been extensively dissected genetically, it is still unknown what forces play a role during migration and neuromast deposition ^34,35^. Therefore, this process of collective cell migration will benefit from an accessible and non-invasive method to estimate forces in a dynamical tissue. Two characteristics of this migratory primordium make it well-suited for applying ForSys: the tissue as a whole is migrating through the lateral line, and its membranes have a curved shape. We showed that ForSys can detect cell division and rosette formation. ForSys will be useful for testing various hypotheses about tissue mechanics in other dynamic cell systems, for instance, during tissue repair and organ regeneration.

We also applied ForSys to address the still mysterious process of planar cell inversion, during which sibling cells rotate around their centroid after the mitotic division of their progenitor ^30^. We discovered that homotypic contacts between rotating cells have the most stress, whereas the contacts between the rotating pair have lower stress than the contacts of each hair cell with its neighbors. This strongly suggests that adhesion dynamics during rotation are based on a strong homotypic interaction of the sibling cells and a weak heterotypic interaction with the surrounding cells, enabling contact exchange during the inversion ^30^.

ForSys provides a versatile and noninvasive tool for studying spatiotemporal patterns of mechanical stresses during tissue morphogenesis in vivo. This software makes stress predictions that can guide researchers in conducting further experiments, which can significantly contribute to understanding the mechanisms involved in development and regeneration. ForSys was built as open-source software in Python, thus allowing the community to participate in its development and maintenance. In our eyes, an interesting future perspective will be to extend the software to tissues in non-equilibrium conditions and adapt the method to operate within a 3D geometry to generate 4D mechanical stress inference.

## Materials and Methods

### SF9 myosin II sensor intensity measurements

Images of stage 16 to stage 20 Xenopus embryos expressing the SF9-3xGFP myosin II sensor were acquired in using a 3i spinning disk microscope with a Plan-Apochromat ×63 oil objective (N.A. = 1.4) mounted on an inverted Zeiss Axio Observer Z1 microscope (Marianas Imaging Workstation [3i— Intelligent Imaging Innovations]), equipped with a CSU-X1 spinning disk confocal head (Yokogawa) and an iXon Ultra 888 EM-CCD camera (Andor Technology). From these images we obtained maximum intensity Z projections, which were then used to extract myosin intensity values of the epithelial junctions. For all vertices constituting a membrane in the segmentation, smoothed intensity values were first obtained by taking the median over first neighbors. The intensity value for each membrane is then defined as the mean of smoothed intensities at each of its constituting vertices. Then, to allow comparison with the inferred stresses, these values were normalized to a mean value of one for each embryo.

### Zebrafish primordium migration experiments

Zebrafish carrying the Tg[−8.0cldnb:Lyn-EGFP]^36^ were kept under standard conditions at 28.5°C. At 40-48 hours post-fertilization, larvae were anesthetized with MS222 and mounted in 0.8% low-melting point agarose on a glass-bottom petri dish. Larvae were imaged in a custom-built Zeiss inverted spinning disk confocal microscope. 16 slices Z stacks of the migrating primordium II were acquired every two minutes with a 63X objective. Subsequently, one z-slice was manually selected from each frame, and the membrane image was segmented using Tissue Analyzer ^37^. The image segmentations were used for ForSys predictions, and the cells’ centroids’ X and Y coordinates, the time point (frame number), and the cell pressures were exported. The probability density function of the cell position along the anteroposterior axis was estimated via a Gaussian kernel in the R statistical software. The value of cell pressure was used as a weight in the density estimation. From this density curve, local maxima were determined through the second derivative.

### The conceptual model behind ForSys

ForSys uses microscopy images as input to estimate the mechanical state of the tissue. The software extracts vertices, edges, and cells from the segmentation, which can be achieved through different software (see, for example, ^8,22,38^). Although most vertices separate two edges, a number of them connect three or more edges and are central for stress inference. We call these pivot vertices or junctions. ForSys calculates the mechanical stresses operating on each edge while assuming mechanical equilibrium in each vertex. Conveniently, this creates a system of equations representing the geometrical state of the tissue ^10,12^. One equation per coordinate is built from every pivot vertex using force balance at the junction.

In the dynamic modality, ForSys assumes that each vertex is in an overdamped regime, where a viscous damping force proportional to the velocity balances the mechanical stresses at that vertex. This creates a non-homogeneous system of equations where the inhomogeneity is proportional to the vertex velocity. In both Static and Dynamic modalities, the resulting system of equations is solved through a Least Squares minimization with the constraint that the average tension equals one (see “Solving the system of equations” in Materials and Methods for more details).

Finally, ForSys uses the stress inferred as an input to estimate cellular pressure within the tissue. For this, a Young-Laplace equation is built at each cell-cell membrane, and the corresponding system is solved similarly to the stresses. However, this requires that the mean pressure of the system is equal to zero (see Section Statical stress inference for details). ForSys renders intercellular stresses as a color code of the cellular outlines, specifically at the edges. Similarly, intracellular pressures are depicted in a color code within the cytoplasmic area of the cells. Moreover, the numerical values of the inference and other observables are easily exportable, facilitating further analysis of the mechanical state of the tissue.

### seAPIpy: generation of *in silico* tissues to validate ForSys

To validate the accuracy of ForSys, we compared the intercellular mechanical stresses inferred by the software with a ground truth distribution of stresses within the tissue. To establish the ground truth, we employed a cell-based computational model to simulate tissues with known intracellular pressures and intercellular stress patterns. Specifically, we employed the vertex model, which is particularly suitable for mechanically evolving epithelial tissues ^39,40^. For the implementation of the vertex model, we utilized Surface Evolver software ^32^. To facilitate the integration and streamline the simulation process, we developed a Python-based software called seapipy ^33^. This open-source computational tool enables Python scripting to generate the desired initial tissue conditions and simulate them using a vertex model implemented in Surface Evolver. seAPIpy generates a Voronoi tessellation with a given geometry as a starting configuration and assigns initial stresses to the edges (Extended data figure 1). Through seAPIpy functions, the user may add Surface Evolver commands to create the desired conditions for evolution and generate the Surface Evolver-compatible file.

By leveraging both ForSys and the capabilities of seAPIpy software, we implemented four conditions as examples that were later used to test ForSys stress inference in both its Static and Dynamic modalities. The first two conditions induce a furrow formation on vertical and horizontal strips, respectively. In the third condition, a central zone of elevated stress is introduced, which diminishes radially. Lastly, a fourth condition assigns five different random stresses to edges, following a uniform distribution, with a 50 % spread in stress values. Each condition underwent twenty-five repetitions. These simulations served as the ground truths for validating ForSys *in silico*, as shown in the following two sections (Fig. 2 A, 3 A, and Extended data figure 2).

We generated four examples to validate our software *in silico*. In all four cases, tissues evolve until a time zero is defined. The stresses are modified according to a prescribed condition, and the tissue evolves for shorter periods while it relaxes.

We generated the initial condition in each example by creating a Voronoi tesselation from N = 64 points in a rectangular grid. Each point in the grid is moved with a Gaussian noise centered at zero. Initial cell target areas are randomly assigned as A = 450 ±5 (mean ± std) from a normal distribution. The initial stress of each edge is also taken from a normal distribution centered at 1 with a standard deviation of 0.1. From this state, the tissue evolved through several rounds of vertex averaging and T1 swaps with varying scales.

We defined time as the number of steps elapsed, times the scale (*t = n Δt*), and call it Surface Evolver Time (SET). The first time point is generated after 3875 SET, after which the tissue is evolved for an additional 125 SET. At this point, membrane stresses are changed according to each condition, and each simulation snapshot is saved every 0.25 SET.

In the conditions corresponding to the horizontal and vertical furrows, the new tensions are generated by summing the value corresponding to the position of the center of an edge in the probability density function (PDF) of a normal distribution to the initial randomized value. The normal distribution has its maximum at the centroid of the tissue and a standard deviation of ∼2 cellular radii. Vertical furrows have the PDF on the y-axis, and horizontal furrows on the x-axis.

Similarly, the circular furrow uses the distance of the edge’s center to the tissue’s centroid to calculate the new stress. Finally, in the condition corresponding to the “random examples”, tensions are assigned from normal distributions with a 50% spread around five possible values (1, 1.1, 1.2, 1.3, 1.5) chosen uniformly.

Therefore, seapipy facilitates testing multiple *in silico* examples and has an easy integration into the analysis pipeline. seapipy offers advantages over an existing package (python-evolver) ^41^ because it incorporates Surface Evolver syntax directly into the Python code, eliminating the need to write Surface Evolver commands manually into the input files. seapipy allows for a systematic and straightforward generation of *in silico* ground truths, enabling a better exploration of strengths and limitations in stress inference tools.

### Statical stress inference

We assume a 2D tissue with C cells, representing each cell as a polygon. The system consists of *V* vertices and *E* edges in total. Each edge is composed of two vertices. We define pivot vertices as those that correspond to junctions between three cells or are at the border of the tissue. This method can be applied to junctions shared by more than three cells but at the risk of losing stability in the underlying model ^42^. All vertices between two pivots are regarded as virtual, and only pivot vertices are used to compute stresses. We then use Newton’s second law and assume mechanical equilibrium to assert that the sum of forces at each pivot vertex equals zero. We can calculate the force acting on each vertex as a sum of the contributions of the forces along the edges connected to it. Mathematically, the force at each pivot vertex will have an equation in the form

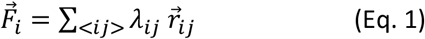

Where *i* and *j* indicate the vertices *i* and *j*, 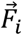 is the force on vertex*i, λ*_*ij*_is the edge force modulus in that edge, and 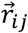 is the versor along the edge starting at vertex*i*. The sum is done over all *j* vertices connected to the vertex *i*. Note that *λ*_*ij*_=*λ*_*ji*_. The directions of the *r*_*ij*_ *versor i*s obtained by fitting a circle to the corresponding membrane, following other authors ^10,12^.

Applying Eq. 1 to all the vertices in the tissue will translate into a homogeneous set of linear equations that have to be solved simultaneously with the edge tensions (*λ*_*ij*_) as unknowns. Hence, we write Eq. 1 and equate it to zero for each system vertex to guarantee that all the forces are balanced. Each of these *V* equations will be written as

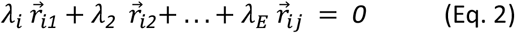

this equation corresponds to the *i*th vertex, and the edge tensions *λ are the* unknowns. Similarly, it is possible to infer the pressures of each cell in the tissue by assembling a system of equations that connects the stress at each membrane with its curvature. The Young-Laplace equation relates these quantities with the pressure difference between two neighboring cells. Symbolically,

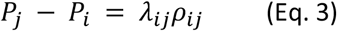

Where *P*_i_ is the pressure of cell *i, λ*_*ij*_ is the stress of the membrane shared by cells *i* and *j*, and *ρ*_*ij*_ is the curvature of the shared membrane. This leads to a system of *E* equations, one per edge, and *C* unknowns.

### The dynamic inference case

The static inference algorithm assumes that vertices do not move. To perform stress inference in a dynamic tissue where all vertices are moving, we modified the static algorithm to include vertex movement. If the system has a low Reynolds number, viscous forces dominate the dynamics over inertial components; Eq. 1 can be modified, assuming a constant viscosity throughout the tissue, to incorporate viscous forces as

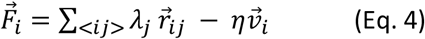

where 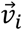 is the velocity of vertex *i* and *η* the viscosity of the tissue. This would modify the coupled system of equations, which could be rearranged to get the ith vertex

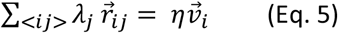

To determine the scales correctly, we proceeded to make Eq. 5 nondimensional. For this, we redefine the stresses by using an unknown reference stress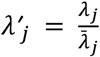. We take this reference stress as the average stress in the system. We used a reference velocity defined as the time average over all the frames of the mean junction velocity

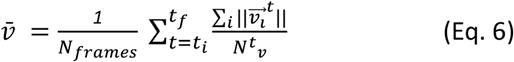

Combining these equations gives a nondimensional expression for the force balance at each junction

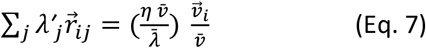

Importantly, this led to the nondimensional parameter 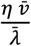. Even though the right-hand side of equation 5 is not a viscosity but rather a damping coefficient, we can interpret it as such in this context. Therefore, as the stress of each membrane represents the elastic forces in the system, this parameter can be interpreted as the relation between the elastic and the viscous forces acting on the system, which is inversely proportional to the Weissenberg number 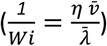.

### Solving the system of equations

In static and dynamic cases, it is necessary to solve a system of linear equations with homogeneous and inhomogeneous conditions, respectively. In both cases, we will turn the system into its matrix form, add a constraint to the unknowns through a Lagrange multiplier, and convert it into a least squares problem. Finally, we will attempt to invert the resulting matrix, and if that is not possible, we will use a numerical algorithm to find the best solution.

Given a two-dimensional tissue with *V* vertices and *E* edges, the system would have 2*V* equations, as each vertex has one equation per dimension and *E* unknowns, one for each edge. Following the method proposed by Brodland *et al*.^10^, the set of equations in Eq. 1 is then translated into matrix form as

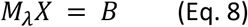

Where *M*_*λ*_ is a 2*V* x *E* matrix with versor coefficients, *X* is the unknowns column matrix of *E* x 1, and *B* is a 2*V* x 1 column matrix with either all zeros under static conditions or the velocity components for each vertex in the dynamical case. To avoid the null solution in the static case, one further condition is added: The mean value for the unknowns, *i*.*e*., *λ*s, is set equal to one, using the equation

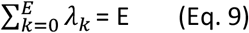

where *E* is the number of edges and *λ*_*k*_ *is the t*ension corresponding to the *k*th edge. In the matrix representation, this entails adding a Lagrange multiplier to the unknowns, a row and column of ones for the tension constraint, and a new row in the *B* matrix. Hence, the equations to be solved have 2*V* + 1 equations and *E* + 1 unknowns. As this system might not always guarantee a solution, we transformed it using least squares. To this end, we apply the transpose matrix *M*^*tr*^_*λ*_ to the equation, giving a new system

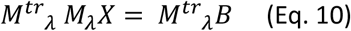

Symbolically,

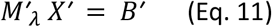

On the other hand, the *B* matrix in Eqs. 6 and 7 for the dynamic case has the corresponding nondimensional velocity component multiplied by the scale parameter (*1/Wi*) described in Eq. 7 in each row. Its final element has the number of edges *E* to enforce the constraint. To quantify the movement present in the tissue, we calculate the 2-norm of the *B* matrix, removing the last row, this vector is referred to as |*v*|_*2*_. Each vertex is tracked through time to obtain the vertex velocity, and the forward velocity is calculated in all but the last step, where the backward expression is used. If a vertex cannot be followed in a frame, i.e., due to significant changes in the tissue shape, it is assigned a null velocity for the frames where it cannot be tracked. Hence, to elucidate the acting forces within the tissue, the software attempts to solve it by inverting the 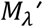 matrix, thus having a solution

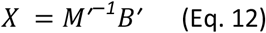

If the system is not invertible, *i*.*e*., *M*’ is singular, or if any of the edge tensions found are negative, a Least Squares algorithm can be used to find the stress values, such as a Non-Negative Least Squares, SciPy’s package or lmfit ^43–45^.

After solving the system, the calculated stresses can be used to infer the pressures of the cells. As seen from the Young-Laplace equation (Eq. 3), pressures are expressed through an inhomogeneous system of linear equations. The left-hand side is a matrix with one column per cell and one row per membrane. Each row has two entries different from zero, one +1 and one −1, representing the difference in pressure at that membrane. The right-hand side consists of a column matrix with the product of each membrane’s stresses and curvatures (*λ*_*ij*_*ρ*_*ij*_). Then, the equations are solved analogously to the stress case using the Least Squares with the constraint that the average pressure must be zero.

### Evaluating goodness of fit

We evaluated the goodness of fit of the inferred data to the ground truth using a tailored saturated score function. This score combines the Pearson correlation coefficient (p), the Mean Absolute Percentage Error (M), and the coefficient of determination (r) as

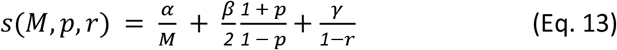

where α, β and γ are free parameters set to one. As this function is unbounded from above, we saturate the score at s = 299.5 for representation purposes in Figure 2, Figure 3 and Extended Data Figure 5. This value comes from an error of 1 %, *i*.*e. s*(*0*.*01, 0*.*99, 0*.*99*) *= 299*.*5*.

### Determination of the scale parameter

We performed a sweep for the correct parametrization of the scale value (1/*Wi*) in the *in silico* examples, from 0 to 0.5. We calculated the score value for each of the five repetitions in the four examples at each time (Extended data figure 5A). We chose the best parameter as the median in each case (Extended data figure 5B). This value coincided with the mode in each case. We used 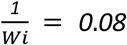 for the x-axis furrow, 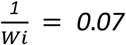 for the y-axis furrow, 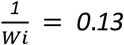 for the circular furrow, and 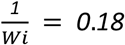 for the random tensions.

### Comparing ForSys with other computational methods

We tested the similarity of the static implementation of ForSys with two other established software: CellFIT^10^ and DLITE^12^. To this end, we applied the DLITE python package to solve the four *in silico* examples used throughout this work, taking advantage of its CellFIT modality. We found that the coefficient of determination is almost equal among the methods (Extended data Fig. 3A) and that the stress distributions emerging from the solution are roughly identical (Extended data Fig. 3B). Moreover, the coefficient of determination is similar for the accumulated data of all repetitions for each example at the last simulated frame (Extended data Fig. 3C).

Moreover, we generated an artificial normal distribution to measure the relative differences with a first moment of 1 and a second moment equal to 0.2. We calculated the Wasserstein Distance between the *in silico* distributions and the normal generated randomly. Given two distributions, X and Y, the Wasserstein Distance is zero if and only if the two distributions are equal. The distance between two distributions can be arbitrarily large for increasingly different shapes.

The Wasserstein Distance is almost zero in all cases, indicating that the distributions gathered from the three inference methods are similar. To compare its similarity, we used an artificially generated normal distribution. Using this metric, we found that the methods among themselves are ∼30 times closer in the x-furrow and y-furrow, 10 times closer for the circular case, and ∼5 times closer in the random densities example than to the normal distribution.

### Statistical estimators

To compare distributions, the Mann-Whitney *U* test was used with different alternative hypotheses, depending on whether we tested for stochastic ordering or whether distributions are different. In all *in silico* cases, the number of samples is twenty-five, which is the number of repetitions per condition. The Pearson correlation coefficient (*R*) was used when we evaluated correlations. The number of samples in each case is indicated when reporting the *p*-value.

## Data availability

All relevant data and materials will be made available upon request.

## Code availability

The seapipy^33^ codebase is available on Github at https://github.com/borgesaugusto/seapipy. ForSys^46^ is available on GitHub https://github.com/borgesaugusto/forsys.

## Acknowledgments

The authors thank Luis Morelli, Fabian Rost, Nicolas Aldecoa, Alice Descoeudres and all the members of the Chara laboratory for valuable comments and suggestions. A.B. was funded by the BMBF 01GQ1904 grant. A.C. was funded by a Doctoral fellowship from CONICET, Argentina. J.S. acknowledges the support of the Novo Nordisk Foundation (grant number NNF19OC0056962) and Leo Foundation (grant number LF-OC-19-000219). The Novo Nordisk Foundation Center for Stem Cell Medicine (reNEW) is supported by a Novo Nordisk Foundation grant number NNF21CC0073729. O.C. was funded by CONICET and Fondo para la Investigación Científica y Tecnológica (PICT-2019-03828).

## Author Contributions

A.B. developed the ForSys code, simulated the in silico results, analyzed all the inference results and wrote the manuscript. A.C. contributed to the development of the ForSys code and analyzed all the inference results. G.V. generated the Xenopus images and edited the manuscript. J.M.-R. acquired the zebrafish microscopy images and analyzed the corresponding results. J.S. supervised the Xenopus project and edited the manuscript. H.L.S. supervised the Zebrafish project and edited the manuscript. O.C. conceived the project, contributed to the ForSys code, analyzed all the inference results, supervised A.B. and A.C as well as wrote the manuscript.

## Extended data figures Legends

**Extended Data Figure 1.**
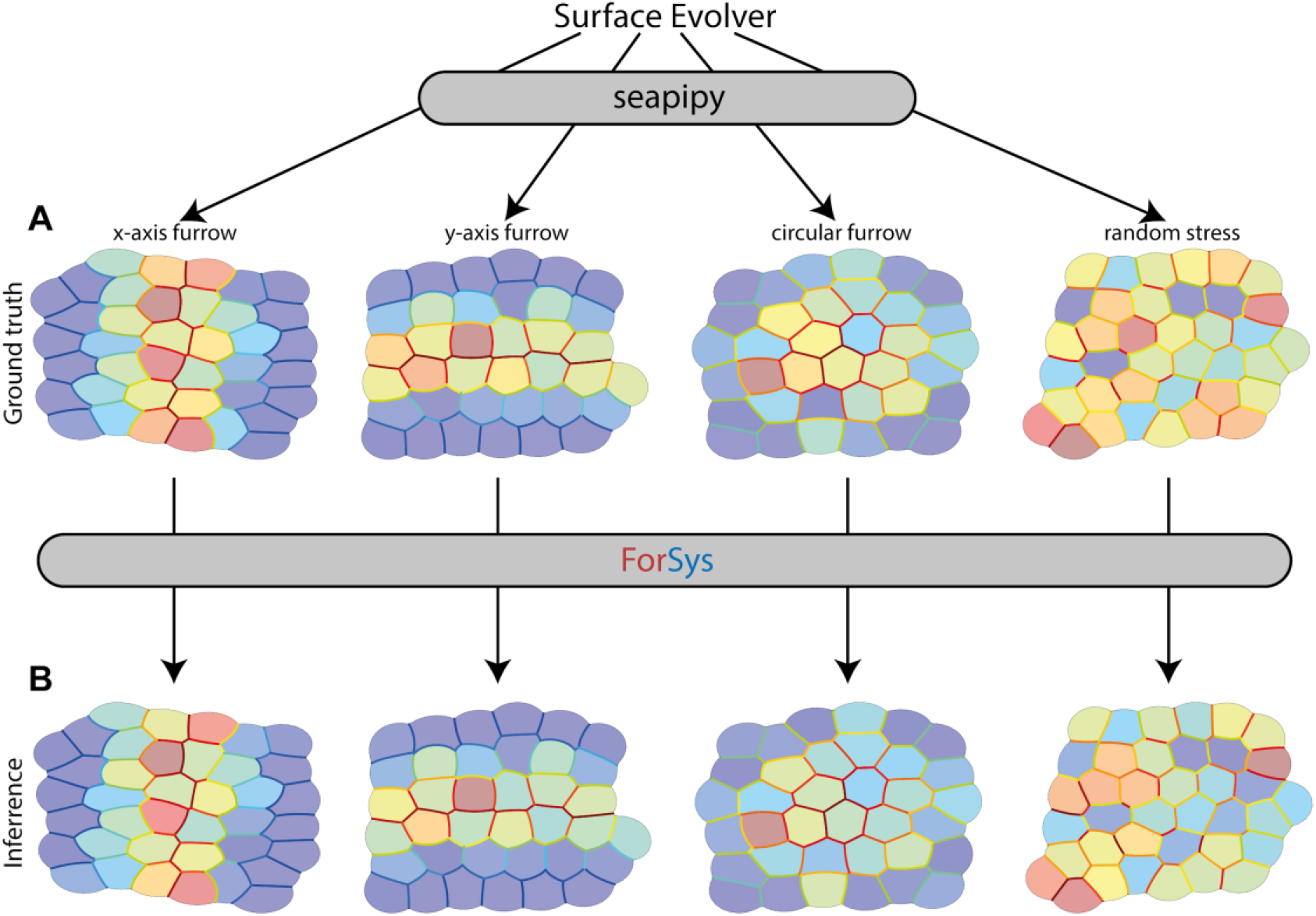
ForSys pipeline for validation. We used different conditions to generate example tissues with varied stresses and pressures. All examples are created through the seapipy package, which uses Surface Evolver as a backend **(A)**. Then, ForSys is applied in any of its modalities to infer the stresses and pressures **(B)**.

**Extended Data Figure 2.**
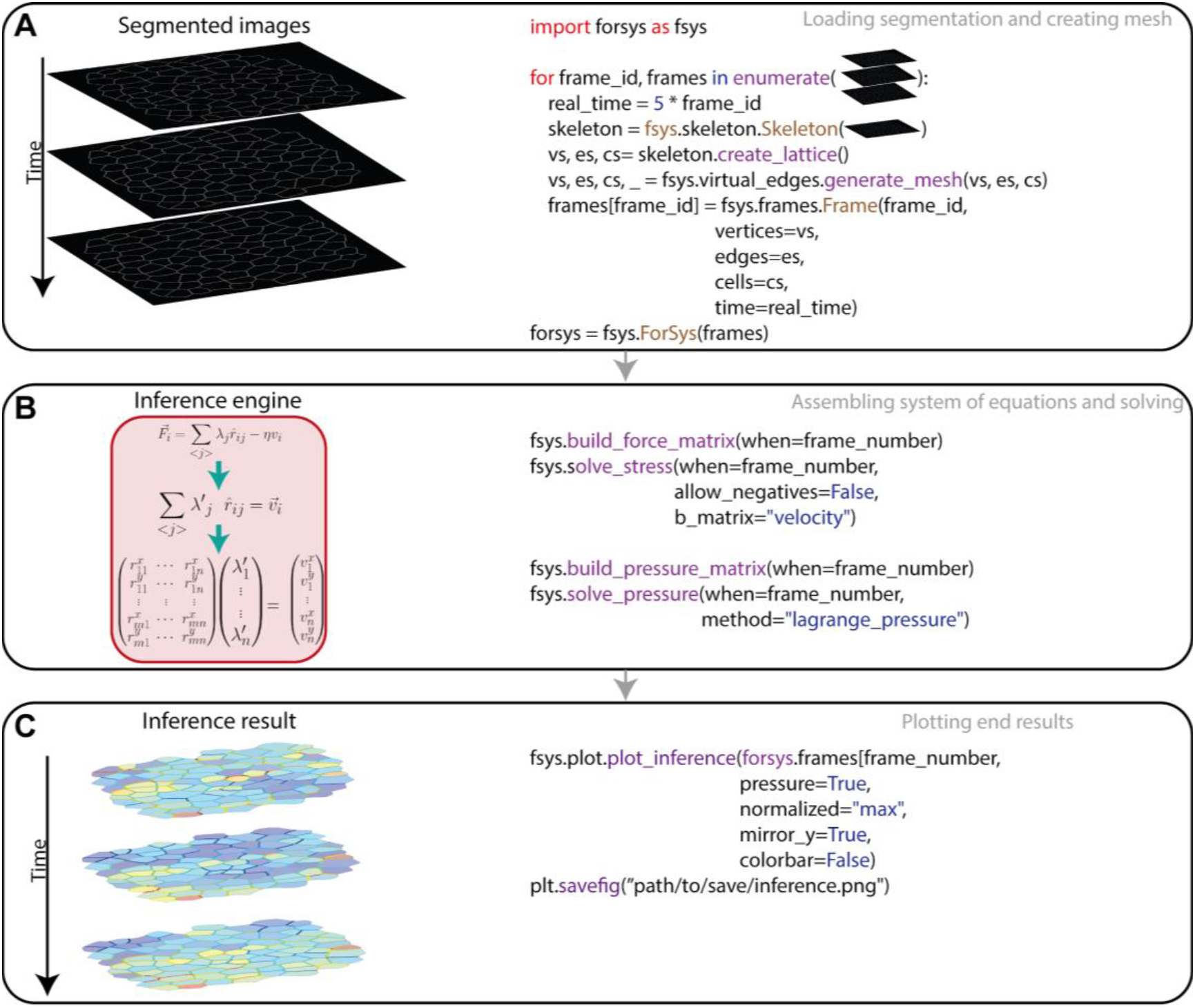
ForSys general implementation. Forsys uses skeletonized images as input, which are read with the Skeleton() module of the software. Frames are collected in a dictionary and then passed to the ForSys() class to generate the corresponding mesh **(A)**. Then, the matrices for the stresses and the pressures are built through different modules and solved individually **(B)**. Stresses must be previously calculated to infer the pressures due to their dependence on the membrane stress. Finally, the inferred stresses and pressures are exported through the *plot_inference*() methods **(C)**.

**Extended Data Figure 3.**
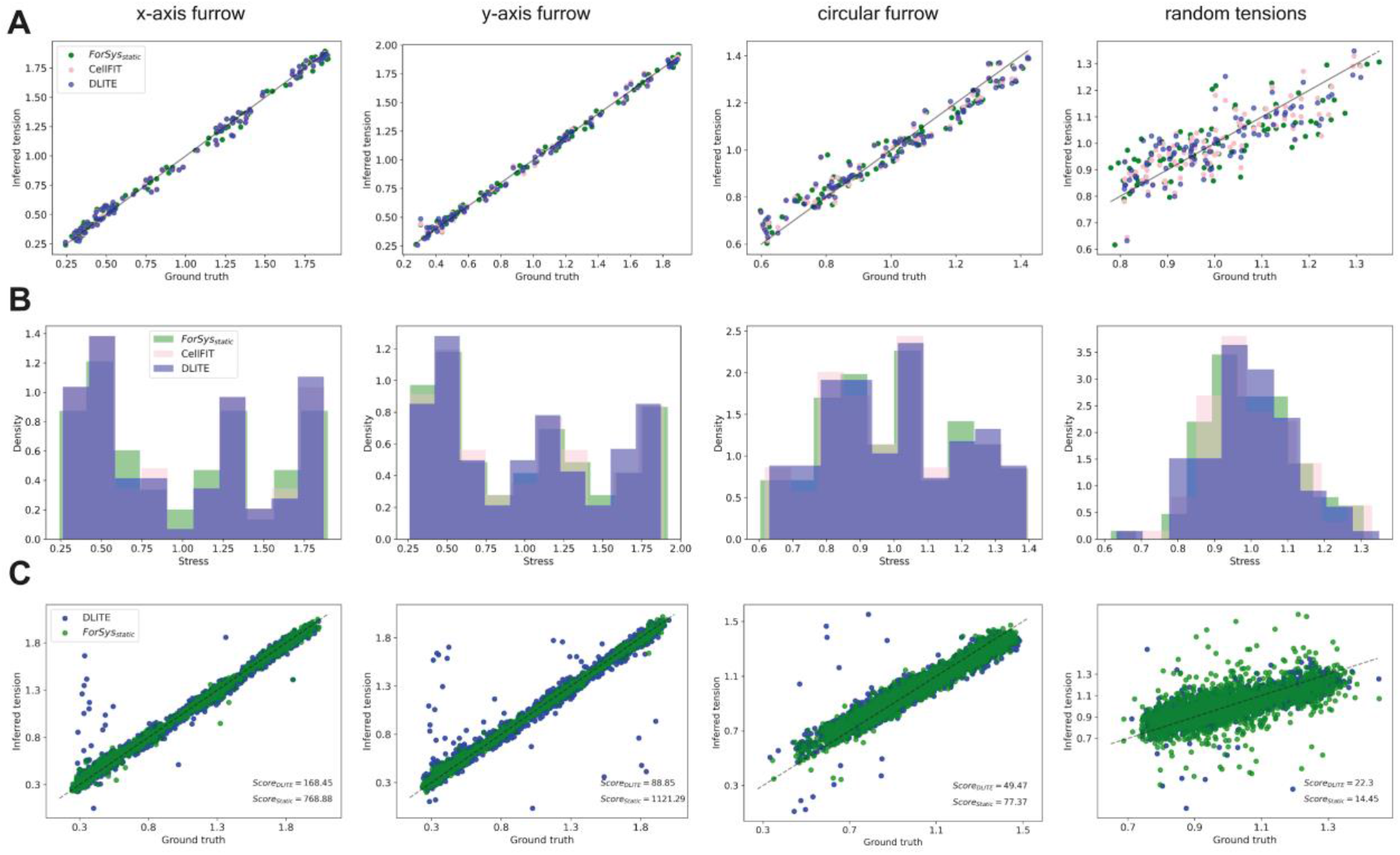
Comparison between staticForSys and other force inference methods. We tested whether the static implementation of ForSys differed from the values of DLITE and CellFIT. Each column represents one of the examples. We show that the inferred stress versus the ground truth follows the y=x line, plotted as a solid black line as a visual aid, for the three methods at the last simulated frame **(A)**. Moreover, the distribution of stresses of all methods has similar behaviors in the histograms **(B)**. Both panels **(A)** and **(B)** are for a selected representative simulation. Then, the result for all inferred tensions versus ground truth repetitions is shown for each condition at the last simulated frame. The black dashed line is the y=x line and is a visual aid. The score function’s values are in the lower right corner of each plot **(C)**. ForSys, in its static modality, has better results in the three first examples and comparable results in the random tension case.

**Extended Data Figure 4.**
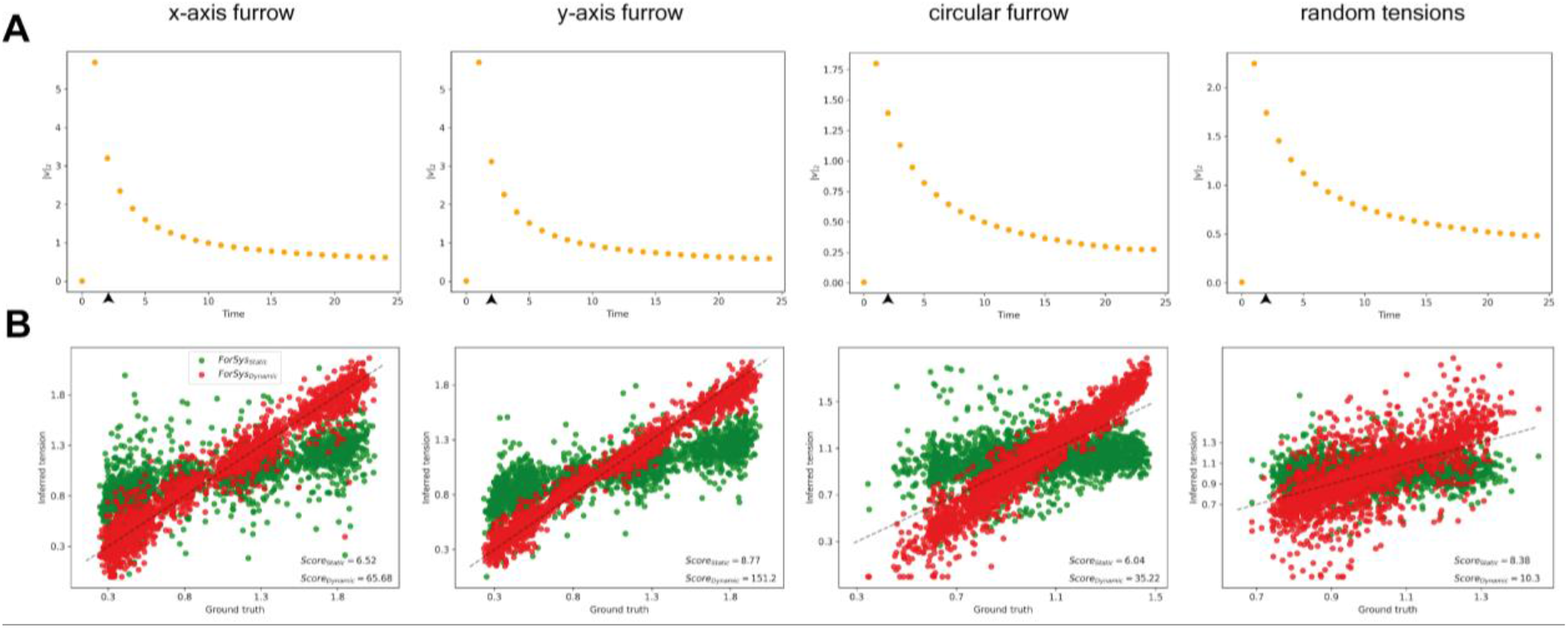
Forsys dynamic at each example. Each column groups plots corresponding to the same prescribed conditions. **(A)** The evolution of tissue movement for each example is shown. The orange scatter dots are the mean of the 2-norm of the velocities vector, with the uncertainty being one standard deviation. In all cases, the values are derived from each example’s repetitions. **(B)** The inferred tension versus ground truth is plotted for all examples. Dynamical results are plotted in red, while static ones are in green. The y = x is plotted as a visual aid as a dashed black line. The score function values are in the lower right corner of each plot. In every case ForSys in its Dynamic modality gives a better score than its static counterpart.

**Extended Data Figure 5.**
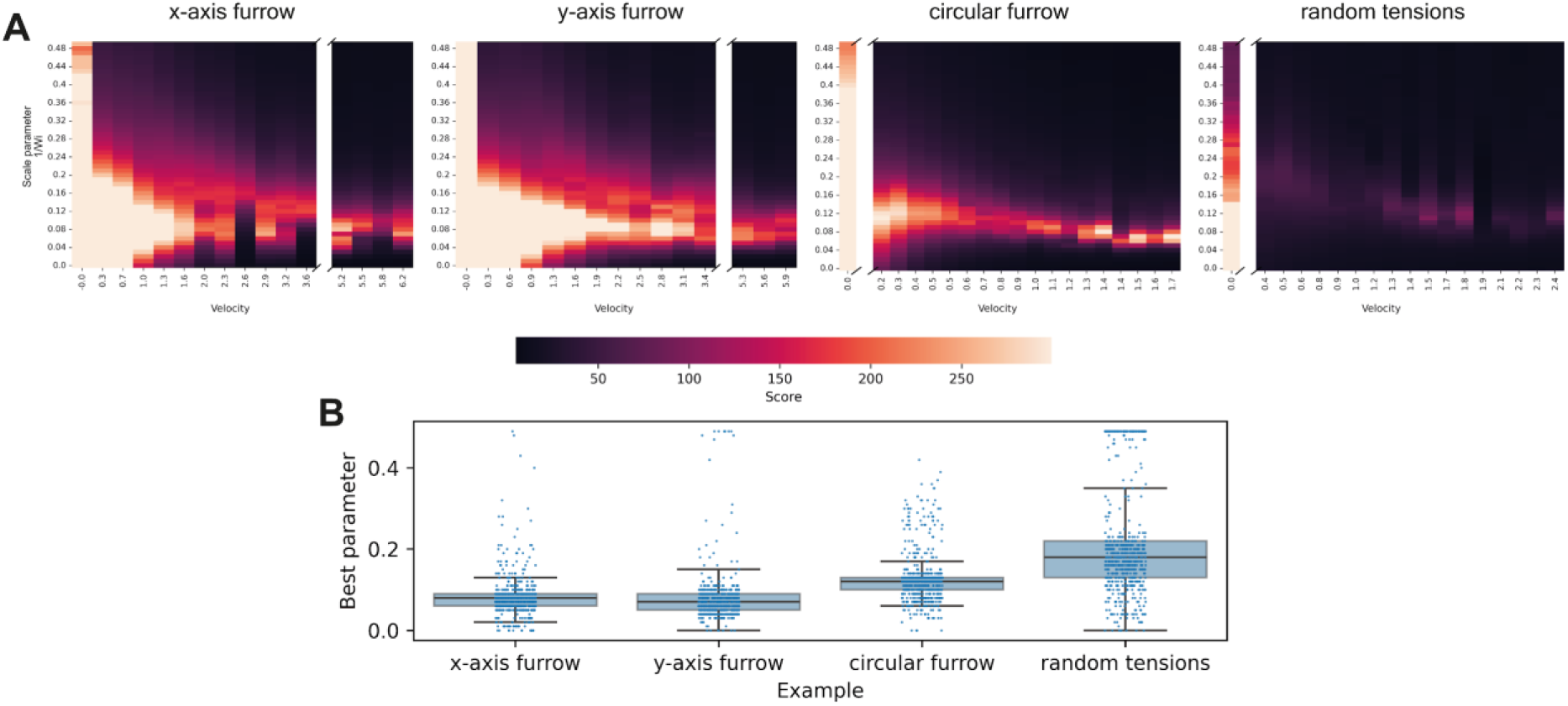
Scale parameter exploration. **(A)** Heatmaps show the saturated score function values for the sweep of the scale parameter proportional to the inverse of the Weissenberg number (*Wi*)^*−1*^ for each example and all 25 repetitions as a function of the velocity of the frame. This velocity is defined as the norm of the vector containing the velocities in the x and y direction of each tracked junction in the tissue. The score function is calculated as described in Materials and Methods, with saturation at the corresponding value *s*(*0*.*01, 0*.*99, 0*.*99*). **(B)** Shows the boxplots for the scale parameter corresponding to the highest score value for each time point and example. The median values of these four distributions are *m*_*x−axis*_ *= 0*.*08; m*_*y−axis*_ *= 0*.*07; m*_*circular*_ *= 0*.*12; m*_*random*_ *= 0*.*18*. The corresponding examples for Figures 2 and 3 use these values as scale parameters.

**Supplementary video 1. ForSys dynamic inference in the migratory primordium**. Top panel shows the time-lapse of the migrating primordium of figure 5 with cell membranes marked with EGFP. The primordium migrates from anterior (left) to posterior (right). The individual frames were segmented and used as input to Forsys. Bottom panel presents the corresponding values of intercelullar stresses and intracellular pressures for each frame as inferred by ForSys. The color of the cell borders and cell area represent the values of the stress and pressure, respectively. Warmer colors indicate higher values and cooler colors indicate lower values.

